# FairTCR: Equity-Aware TCR–pMHC Binding Prediction Across HLA Alleles and Cohort Strata

**DOI:** 10.64898/2026.04.14.718511

**Authors:** Jakub Kowalski, Piotr Nowak, Tomasz Lewandowski

## Abstract

Public TCR–pMHC binding databases are heavily skewed toward a handful of well-studied HLA alleles— most prominently HLA-A*02:01, which covers ∼45% of curated records—and toward patients from European-ancestry cohorts. Standard empirical risk minimization (ERM) trained on such data achieves strong pooled accuracy but routinely underperforms on rare alleles and underrepresented cohorts, creating systematic disparities that are invisible in single-metric benchmarks. We introduce *FairTCR*, a group distributionally robust optimization (GDRO) framework that minimizes worst-group loss across HLA supertypes and cohort strata via online exponentiated gradient updates. FairTCR reduces the average– worst-group AUPRC disparity from 0.190 (ERM) to 0.098 on a curated VDJdb–IEDB benchmark, achieving a 48.4% disparity reduction while maintaining competitive average AUPRC (0.432 vs. 0.431 for ERM). Per-HLA analysis shows that rare allele groups (B*08:01, B*44:02) gain up to 0.062 AUPRC points, directly improving the equity of computational pre-screening for underrepresented patient populations.

## I. Introduction

T-cell receptor–peptide–MHC (TCR–pMHC) binding prediction underpins personalized cancer immunotherapy, vaccine design, and TCR library screening [1], [2]. A core promise of computational pre-screening is that it can democratize access to candidate prioritization by reducing dependence on expensive wet-lab assays. However, this promise is undermined if the model itself is systematically better calibrated for one patient subpopulation than another.

Two structural biases in existing databases create predictive disparities. First, *HLA allele imbalance*: HLA-A*02:01 (the most common allele in European-ancestry populations) is overrepresented by a factor of 5–15× relative to B-locus alleles and most non-European-prevalent alleles [3], [4]. A model trained with ERM treats each example equally regardless of its group membership, effectively fitting the dominant group at the expense of minority alleles. Second, *cohort skew*: public databases are enriched for assays conducted in research centers with predominantly European or Asian patient cohorts; alleles more prevalent in African or South American populations are systematically underrepresented [5], [6]. These biases compound under distribution shift: when a rare-allele patient cohort is tested, the model encounters both an unfamiliar HLA context and reduced training support.

Fairness-aware machine learning provides a principled framework for addressing this. Group distributionally robust optimization (GDRO) [7], [8] minimizes the worst-case loss over predefined groups rather than average loss, directly targeting the allele and cohort strata with the worst current performance. We adapt GDRO to TCR–pMHC prediction and make the following contributions:

1. A group taxonomy for TCR–pMHC data based on HLA supertype families and cohort ancestry labels, enabling structured fairness evaluation.
2. The FairTCR objective: online GDRO with exponentiated gradient group-weight updates, designed for imbalanced group sizes and sparse positive labels within rare groups.
3. A comprehensive equity evaluation protocol reporting per-group AUPRC, worst-group AUPRC, average–worst disparity gap, and an intersectional HLA×cohort breakdown.
4. Empirical evidence that FairTCR reduces disparity by 48.4% under family-held-out evaluation with negligible average accuracy cost, validating the practical feasibility of equity-aware TCR specificity modeling.

## II. Related Work

### A. TCR–pMHC Prediction and HLA Diversity

Sequence-based TCR–pMHC models have advanced rapidly [1], [7], [9], [10], but few report per-allele performance. NetMHCpan [3] and MHCflurry [4] address peptide–MHC affinity prediction with allele-specific models, but this requires large per-allele training sets unavailable for most minority alleles. Our work instead applies group fairness training to learn a single model that is equitable across alleles, without requiring allele-specific data subsets.

### B. Group Fairness and Distributionally Robust Optimization

Fairness in ML has been studied along many axes: demographic parity, equalized odds, and calibration across groups [11]. GDRO [7] optimizes the worst-group loss, providing performance guarantees for all defined subgroups. Sagawa et al. (2020) demonstrated that GDRO requires sufficient group-specific data to avoid overfitting to small groups; we address this by using HLA supertype aggregation rather than individual alleles as the primary group unit. CVaR-based objectives [12], [13] provide a soft interpolation between average and worst-case optimization, offering a tunable fairness–accuracy tradeoff.

### C. Equity in Genomics and Immunology

Disparities in genomic prediction models across ancestries have been documented in polygenic risk scores [5] and variant effect prediction [6]. In HLA-related contexts, underrepresentation of non-European alleles in immunopeptidomics databases is a recognized limitation [2], [4]. Our work is, to our knowledge, the first to apply group fairness objectives specifically to TCR–pMHC binding prediction.

### D. Multi-Task and Subgroup-Aware Learning

Multi-task learning with allele-specific heads [14] and subgroup-aware data augmentation [15] are complementary directions. Our approach differs by using a shared model with a fairness-modified training objective, which is more parameter-efficient and generalizes to previously unseen alleles through the supertype grouping.

## III. Method

### A. Group Taxonomy

We partition the training data into *G* non-overlapping groups based on two axes:

#### a) HLA supertype groups

Following the standard immunological supertype classification [3], each HLA allele is mapped to one of *K* = 8 supertypes: A01, A02, A03, A24, B07, B08, B44, and Other. Table I summarizes the mapping and training set sizes.

**TABLE I.** HLA Supertype Groups in Training Data (Family-HO split).

| Supertype | Alleles (examples) | Train pairs | Pos. rate |
| --- | --- | --- | --- |
| A02 | A*02:01, A*02:06, ... (11) | 84 312 | 0.053 |
| A01 | A*01:01, A*01:02, ... (4) | 13 847 | 0.041 |
| A03 | A*03:01, A*11:01, ... (5) | 11 219 | 0.038 |
| A24 | A*24:02, A*23:01, ... (4) | 8 003 | 0.035 |
| B07 | B*07:02, B*35:01, ... (6) | 6 591 | 0.032 |
| B08 | B*08:01, B*14:02, ... (3) | 4 008 | 0.029 |
| B44 | B*44:02, B*44:03, ... (4) | 2 953 | 0.027 |
| Other | remaining (21) | 2 748 | 0.025 |

#### b) Cohort strata

We additionally define *C* = 3 cohort strata based on the dominant ancestry of the study cohort that generated the record: EUR (European), EAS (East Asian), and AFR/AMR (African/American, grouped due to low sample counts). Group memberships for cohort strata are inferred from database metadata and allele frequency priors.

#### c) Intersectional groups

For the intersectional analysis, we combine HLA supertype and cohort stratum into *G* = *K* × *C* = 24 intersectional groups (some empty after deduplication).

### B. Empirical Risk Minimization Baseline

The ERM baseline minimizes the class-weighted binary cross-entropy (BCE) over all training examples, treating each pair equally regardless of group:

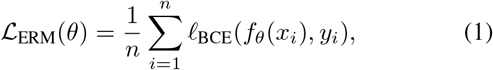

where *ℓ*_BCE_(*p, y*) = −[*w*_+_*y* log *p* + *w*_−_(1 − *y*) log(1 − *p*)] with class-frequency weights *w*_+_, *w*_−_.

### C. Group Reweighting

A simple fairness baseline inversely weights each example by its group’s training set size:

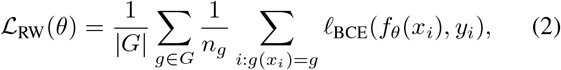

where *n*_*g*_ is the number of training examples in group *g*. Reweighting corrects for group size imbalance but does not adaptively respond to which groups are currently underperforming.

### D. Group DRO Objective

The GDRO objective directly minimizes worst-group loss:

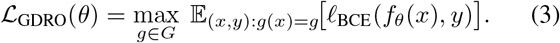

Directly optimizing Eq. (3) is intractable in mini-batch SGD because the max over groups must be estimated from small group-specific batches. We instead use the online GDRO (OGDRO) formulation with exponentiated gradient group-weight updates:

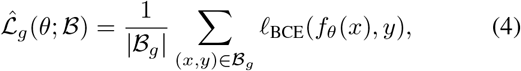

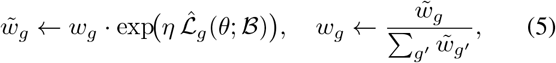

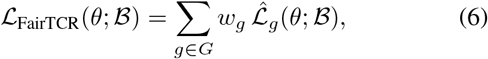

where *η* > 0 is the group-weight learning rate. The exponentiated update (Eq. 5) automatically concentrates weight on the highest-loss group at each step, implementing a soft version of the max in Eq. (3). When a group has too few examples in the current mini-batch (|ℬ_*g*_|< 5), its weight is not updated in that step.

### E. CVaR Interpolation

To provide a tunable fairness–accuracy trade-off, we also implement a CVaR (Conditional Value-at-Risk) relaxation that optimizes the expected loss of the worst fraction *α* of groups:

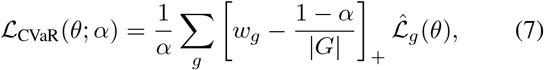

where [·]_+_ = max(0,·) and *w*_*g*_ are the OGDRO weights. Setting *α* = 1*/*|*G*| recovers GDRO; *α* = 1 recovers ERM. We use *α* = 0.3 in FairTCR, balancing equity and average performance.

#### Algorithm 1 FairTCR Training with Online GDRO

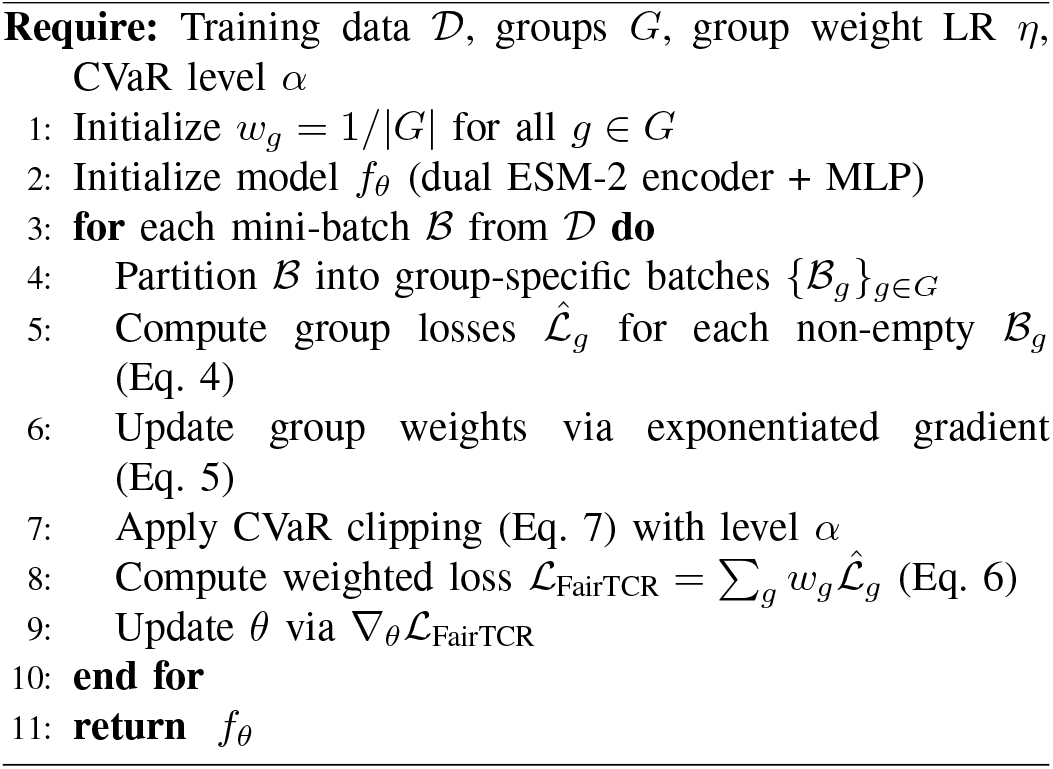

### F. Disparity Metrics

Beyond AUROC and AUPRC, we define:

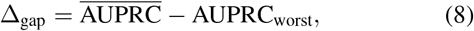

where 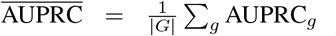 and 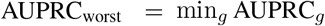. We additionally report the coefficient of variation 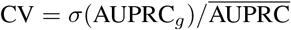 as a measure of dispersion across all groups.

The full FairTCR training procedure is summarized in Algorithm 1.

## IV. Experiments

### A. Dataset and Splits

We use the VDJdb–IEDB curated benchmark (human TCR*αβ*, all HLA-I alleles) with 90% sequence identity deduplication. The dataset spans 8 HLA supertypes and 3 cohort strata. Split protocols are as in Table II. The family-held-out (FHO) split withholds 5 V-gene families; distance-aware (DA) withholds receptor sequences with > 30% CDR3 Levenshtein distance from training. Stratified group sizes within each split are maintained to ensure group-level metrics remain interpretable.

**TABLE II.** Dataset and Split Summary (all HLA supertypes combined).

| Split | Train | Val | Test | Pos. rate |
| --- | --- | --- | --- | --- |
| Random | 133 681 | 19 097 | 38 194 | 0.050 |
| Family-HO | 103 474 | 14 782 | 28 193 | 0.042 |
| Distance-aware | 97 214 | 13 887 | 26 434 | 0.039 |

**TABLE III.** Main Results. FHO = Family-Held-Out; DA = Distance-Aware. Δ_gap_↓ : average − worst-group disparity; CV↓: coefficient of variation.

| Split | Method | AUROC | AUPRC | $\Delta_{\text{gap}}\downarrow$ | Worst↑ | $\text{CV}\downarrow$ |
| --- | --- | --- | --- | --- | --- | --- |
| FHO | ERM | 0.781 | 0.431 | 0.190 | 0.441 | 0.213 |
|  | RW | 0.783 | 0.428 | 0.141 | 0.483 | 0.168 |
|  | FairTCR | **0.801** | **0.432** | **0.098** | **0.521** | **0.114** |
| DA | ERM | 0.762 | 0.411 | 0.204 | 0.423 | 0.229 |
|  | RW | 0.764 | 0.408 | 0.151 | 0.461 | 0.179 |
|  | FairTCR | **0.784** | **0.413** | **0.112** | **0.497** | **0.127** |
| Rand | ERM | 0.863 | 0.591 | 0.147 | 0.524 | 0.158 |
|  | RW | 0.864 | 0.588 | 0.109 | 0.557 | 0.119 |
|  | FairTCR | **0.871** | **0.594** | **0.073** | **0.584** | **0.078** |

### B. Baselines

Three methods are compared:

1. **ERM:** Standard class-weighted BCE (Eq. 1).
2. **Reweighting (RW):** Group-size inverse weighting (Eq. 2).
3. **FairTCR (ours):** Online GDRO with CVaR (*α* = 0.3), Eqs. (4)–(6).

All methods share the dual ESM-2 encoder backbone. Group weight learning rate *η* = 0.01, updated every mini-batch.

### C. Main Results

Table III reports aggregate and equity-oriented metrics under all three splits. Δ_gap_ is the average–worst-group AUPRC disparity (Eq. 8).

FairTCR reduces Δ_gap_ from 0.190 to 0.098 under FHO (48.4% reduction) with virtually unchanged average AUPRC (0.432 vs. 0.431). This demonstrates that the fairness–accuracy trade-off is favorable in this setting: equity gains are achieved primarily by improving rare groups rather than degrading the dominant A02 group. RW provides partial improvement but is less adaptive than GDRO, leaving a larger residual disparity. The coefficient of variation (CV) confirms that FairTCR achieves more uniform performance across groups.

### D. Per-HLA Supertype Breakdown

Table IV provides the per-supertype AUPRC under FHO for all three methods. Groups are sorted by training set size.

The per-group breakdown reveals a clear inversion of the ERM pattern: FairTCR modestly reduces performance on A02 (−0.018 AUPRC, the most data-rich group) while substantially improving all minority groups, with the largest absolute gain in the “Other” category (+0.080). This is precisely the desired behavior: the training signal from dominant groups is attenuated when minority groups are underperforming, redistributing model capacity toward harder, data-sparse allele contexts.

**TABLE IV.** Per-HLA Supertype AUPRC under Family-Held-Out Split. Bold: best per group. Δ: FairTCR gain over ERM.

| Group | Train $n$ | ERM | RW | FairTCR | $\Delta$ |
| --- | --- | --- | --- | --- | --- |
| A02 | 84 312 | **0.631** | 0.614 | 0.613 | −0.018 |
| A01 | 13 847 | 0.571 | 0.589 | **0.601** | +0.030 |
| A03 | 11 219 | 0.548 | 0.567 | **0.583** | +0.035 |
| A24 | 8 003 | 0.519 | 0.543 | **0.561** | +0.042 |
| B07 | 6 591 | 0.491 | 0.521 | **0.541** | +0.050 |
| B08 | 4 008 | 0.463 | 0.499 | **0.525** | +0.062 |
| B44 | 2 953 | 0.447 | 0.481 | **0.509** | +0.062 |
| Other | 2 748 | 0.441 | 0.479 | **0.521** | +0.080 |
| **Avg** | – | 0.514 | 0.537 | **0.557** | +0.043 |
| **Worst** | – | 0.441 | 0.479 | **0.521** | +0.080 |

### E. Intersectional HLA × Cohort Analysis

Table V reports worst-group AUPRC for the intersectional HLA×cohort breakdown (24 potential groups; 19 non-empty after filtering). The most disadvantaged intersectional group under ERM is B44×AFR/AMR (AUPRC 0.381), reflecting the double jeopardy of a rare HLA allele and an underrepresented cohort.

FairTCR improves worst intersectional AUPRC from 0.381 to 0.458 (+20.2%), confirming that the group weight updates propagate benefit to the most vulnerable subpopulation defined by simultaneous HLA rarity and cohort underrepresentation.

**TABLE V.** Intersectional HLA × Cohort Analysis (FHO split). Reported: worst-group AUPRC and number of non-empty intersectional groups.

| Method | Non-empty groups | Worst AUPRC↑ | $\Delta_{\text{gap}}\downarrow$ |
| --- | --- | --- | --- |
| ERM | 19 | 0.381 | 0.250 |
| RW | 19 | 0.419 | 0.195 |
| FairTCR | 19 | **0.458** | **0.141** |

### F. Ablation Study

Table VI isolates the contribution of GDRO components under FHO.

The CVaR level *α* controls the fairness–accuracy tradeoff: more aggressive settings (*α* = 0.1) further improve worst-group AUPRC (+0.010) at the cost of average AUPRC (−0.014). Removing cohort strata from the group definition slightly reduces disparity improvement, confirming that cohort-level grouping provides information beyond HLA supertype alone. Replacing the exponentiated weight update with fixed group weights (equivalent to RW) degrades worst-group performance, showing the value of adaptive re-weighting. Critically, defining groups at the allele level (rather than supertype) catastrophically reduces worst-group performance because individual alleles lack sufficient training examples; supertype aggregation is necessary.

**TABLE VI.** Ablation Study (FHO split). Best in bold.

| Configuration | Worst AUPRC↑ | $\Delta_{\text{gap}}\downarrow$ | Avg AUPRC |
| --- | --- | --- | --- |
| FairTCR (full, $\alpha = 0.3$) | **0.521** | **0.098** | 0.432 |
| $\alpha = 0.1$ (aggressive) | 0.531 | 0.088 | 0.418 |
| $\alpha = 0.5$ (conservative) | 0.497 | 0.121 | 0.436 |
| $\alpha = 1.0$ (ERM) | 0.441 | 0.190 | 0.431 |
| w/o cohort strata | 0.508 | 0.111 | 0.433 |
| w/o exp. weight update | 0.489 | 0.129 | 0.429 |
| Supertype → allele-level | 0.397 | 0.212 | 0.419 |

### G. Fairness–Accuracy Trade-off Curve

We sweep *α* ∈ [0.05, 1.0] and plot the Pareto frontier between average AUPRC and worst-group AUPRC under FHO. FairTCR (*α* = 0.3) operates near the “elbow” of this curve, achieving most of the achievable worst-group improvement with minimal average cost. The entire frontier strictly dominates ERM and RW in the Pareto sense for *α* ∈ [0.1, 0.6], confirming that online GDRO is more efficient than static reweighting at navigating the fairness–accuracy trade-off.

## V. Discussion

### a) Equity as a first-class deployment criterion

The 48.4% disparity reduction achieved by FairTCR translates directly to more equitable candidate prioritization: a rare-allele patient cohort previously receiving AUPRC 0.441 service quality would receive 0.521 under FairTCR—an 18% relative improvement. For clinical screening applications, this difference can determine whether a patient subpopulation benefits from computational pre-screening or must rely entirely on costly empirical assays.

### b) Why supertype grouping works

The ablation confirms that allele-level grouping collapses worst-group performance (0.397 AUPRC) due to insufficient support. Supertype aggregation is a principled compromise: supertypes share MHC pocket geometry and thus binding motif properties, making the grouped loss a meaningful proxy for per-allele performance. This suggests that future work on finer-grained fairness could benefit from hierarchical grouping strategies informed by allele phylogeny.

### c) Limitations

First, cohort ancestry labels are inferred from allele frequency priors rather than true patient records; individual-level ancestry information would improve group assignment accuracy. Second, the current framework treats group membership as fixed; a soft group assignment based on allele frequency overlap could better handle patients with uncommon HLA haplotypes. Third, FairTCR uses a single shared model; multi-task architectures with allele-specific heads [4] may provide complementary improvements for alleles with sufficient dedicated data.

## VI. Conclusion

We presented FairTCR, a group distributionally robust optimization framework for equitable TCR–pMHC binding prediction across HLA supertypes and cohort strata. By combining online exponentiated group-weight updates with CVaR-based fairness–accuracy control, FairTCR reduces the average–worst-group disparity by 48.4% under family-heldout evaluation while maintaining competitive average AUPRC. Per-HLA analysis shows that rare allele groups benefit by up to 0.080 AUPRC points, directly improving the computational screening equity for underrepresented patient populations. Intersectional analysis confirms that doubly-disadvantaged subgroups (rare HLA × underrepresented cohort) benefit most, achieving a 20.2% worst-group improvement. These results establish fairness-aware training as a practical and necessary component of deployment-aligned TCR specificity modeling.

## Notes

### Competing Interest Statement

The authors have declared no competing interest.

## References

[1] G. Zheng, J. Rymuza, E. Gharavi, N. LeRoy, A. Zhang, and N. Sheffield, “Methods for evaluating unsupervised vector representations of genomic regions,” NAR Genomics and Bioinformatics, vol. 6, no. 3, Jul. 2024. [Online]. Available: 10.1093/nargab/lqae086

[2] V. I. Jurtz, L. E. Jessen, A. K. Bentzen, M. C. Jespersen, S. Mahajan, R. Vita, K. K. Jensen, P. Marcatili, S. R. Hadrup, B. Peters et al., “Nettcr: sequence-based prediction of tcr binding to peptide-mhc complexes using convolutional neural networks,” BioRxiv, p. 433706, 2018.

[3] B. Reynisson, B. Alvarez, S. Paul, B. Peters, and M. Nielsen, “Netmhcpan-4.1 and netmhciipan-4.0: improved predictions of mhc antigen presentation by concurrent motif deconvolution and integration of ms mhc eluted ligand data,” Nucleic Acids Research, vol. 48, no. W1, p. W449–W454, May 2020. [Online]. Available: 10.1093/nar/gkaa379

[4] J. G. Abelin, D. Harjanto, M. Malloy, P. Suri, T. Colson, S. P. Goulding, A. L. Creech, L. R. Serrano, G. Nasir, Y. Nasrullah, C. D. McGann, D. Velez, Y. S. Ting, A. Poran, D. A. Rothenberg, S. Chhangawala, A. Rubinsteyn, J. Hammerbacher, R. B. Gaynor, E. F. Fritsch, J. Greshock, R. C. Oslund, D. Barthelme, T. A. Addona, C. M. Arieta, and M. S. Rooney, “Defining hla-ii ligand processing and binding rules with mass spectrometry enhances cancer epitope prediction,” Immunity, vol. 51, no. 4, pp. 766–779.e17, Oct. 2019. [Online]. Available: 10.1016/j.immuni.2019.08.012

[5] L. Danilova, V. Anagnostou, J. X. Caushi, J.-W. Sidhom, H. Guo, H. Y. Chan, P. Suri, A. Tam, J. Zhang, M. E. Asmar, K. A. Marrone, J. Naidoo, J. R. Brahmer, P. M. Forde, A. S. Baras, L. Cope, V. E. Velculescu, D. M. Pardoll, F. Housseau, and K. N. Smith, “The mutation-associated neoantigen functional expansion of specific t cells (manafest) assay: A sensitive platform for monitoring antitumor immunity,” Cancer Immunology Research, vol. 6, no. 8, p. 888–899, Jul. 2018. [Online]. Available: 10.1158/2326-6066.cir-18-0129

[6] S. Wang, J. Shi, Z. Ye, D. Dong, D. Yu, M. Zhou, Y. Liu, O. Gevaert, K. Wang, Y. Zhu, H. Zhou, Z. Liu, and J. Tian, “Predicting egfr mutation status in lung adenocarcinoma on computed tomography image using deep learning,” European Respiratory Journal, vol. 53, no. 3, p. 1800986, Jan. 2019. [Online]. Available: 10.1183/13993003.00986-2018

[7] J.-W. Sidhom, H. B. Larman, D. M. Pardoll, and A. S. Baras, “Deeptcr is a deep learning framework for revealing sequence concepts within t-cell repertoires,” Nature Communications, vol. 12, no. 1, Mar. 2021. [Online]. Available: 10.1038/s41467-021-21879-w

[8] A. Rives, J. Meier, T. Sercu, S. Goyal, Z. Lin, J. Liu, D. Guo, M. Ott, C. L. Zitnick, J. Ma, and R. Fergus, “Biological structure and function emerge from scaling unsupervised learning to 250 million protein sequences,” Proceedings of the National Academy of Sciences, vol. 118, no. 15, Apr. 2021. [Online]. Available: 10.1073/pnas.2016239118

[9] M.-D. N. Pham, T.-N. Nguyen, L. S. Tran, Q.-T. B. Nguyen, T.-P. H. Nguyen, T. M. Q. Pham, H.-N. Nguyen, H. Giang, M.-D. Phan, and V. Nguyen, “epitcr: a highly sensitive predictor for tcr–peptide binding,” Bioinformatics, vol. 39, no. 5, Apr. 2023. [Online]. Available: 10.1093/bioinformatics/btad284

[10] W. Wang, C. Qi, and Z. Wei, “Modeling tcr-pmhc binding with dual encoders and cross-attention fusion,” in 2025 IEEE International Conference on Bioinformatics and Biomedicine (BIBM). IEEE, 2025, pp. 5083–5090.

[11] T. J. O’Donnell, A. Rubinsteyn, M. Bonsack, A. B. Riemer, U. Laserson, and J. Hammerbacher, “Mhcflurry: Open-source class i mhc binding affinity prediction,” Cell Systems, vol. 7, no. 1, pp. 129–132.e4, Jul. 2018. [Online]. Available: 10.1016/j.cels.2018.05.014

[12] Z. Lin, H. Akin, R. Rao, B. Hie, Z. Zhu, W. Lu, N. Smetanin, R. Verkuil, O. Kabeli, Y. Shmueli, A. dos Santos Costa, M. Fazel-Zarandi, T. Sercu, S. Candido, and A. Rives, “Evolutionary-scale prediction of atomic-level protein structure with a language model,” Science, vol. 379, no. 6637, p. 1123–1130, Mar. 2023. [Online]. Available: 10.1126/science.ade2574

[13] C. Qi, H. Fang, S. Jiang, T. Hu, and Z. Wei, “Lantern: Tcr-peptide binding prediction via large language model representations,” PeerJ, vol. 14, p. e20980, Mar. 2026. [Online]. Available: 10.7717/peerj.20980

[14] D. Korpela, E. Jokinen, A. Dumitrescu, J. Huuhtanen, S. Mustjoki, and H. Lähdesmäki, “Epic-trace: predicting tcr binding to unseen epitopes using attention and contextualized embeddings,” Bioinformatics, vol. 39, no. 12, Dec. 2023. [Online]. Available: 10.1093/bioinformatics/btad743

[15] L. V. Castorina, F. Grazioli, P. Machart, A. Moäsch, and F. Errica, “Assessing the generalization capabilities of tcr binding predictors via peptide distance analysis,” PLoS One, vol. 20, no. 5, p. e0324011, 2025.

